# Talking avatars can differentially modulate cortical speech tracking in the high and in the low delta band

**DOI:** 10.64898/2026.01.07.695461

**Authors:** Jasmin Riegel, Alina Schüller, Constantin Jehn, Alexander Wißmann, Steffen Zeiler, Dorothea Kolossa, Tobias Reichenbach

## Abstract

In noisy listening environments, visual cues from a speaker’s face can significantly boost speech compre-hension. The underlying audiovisual integration in the brain involves neural tracking of audiovisual speech features. Moreover, lip reading in silence is associated with tracking of the speech envelope in the low-delta frequency band (0.5 *−* 1 Hz). Recently, digital avatars have emerged that can support speech comprehension. Yet, it remains unclear how the human brain integrates such artificial visual signals with natural speech. Here, we employed magnetoencephalography (MEG) to measure the neural response to a natural video, an avatar generated by deep neural networks, and a degraded video serving as a control. We demonstrate that the avatar can enhance speech-in-noise comprehension to a similar degree as the degraded video, although less than the natural video. We further identify a late response at 600 ms in the neural tracking of the auditory cortex in the high delta band (1 *−* 4 Hz) that predicts audiovisual speech comprehension. In contrast, we found that neural tracking in the low delta band is related to silent lip-reading performance. Importantly, the tracking in the low delta band evoked by the avatars is much weaker and occurs earlier than that elicited by the other audiovisual stimuli. Neural tracking in the theta band (4 *−* 8 Hz) is not involved in audiovisual integration. Our results show that the low delta band and the high delta band play clearly distinct roles in visual-only and audiovisual speech processing, and suggest potential avenues for further boosting the abilities of avatars to support speech comprehension.

**Significance Statement:** Understanding a conversational partner is essential for everyday communication. Yet, many people — due to aging or other factors — struggle to follow speech in noisy environments. Seeing the speaker’s face can greatly enhance speech comprehension, but visual cues are often unavailable, such as during public announcements or telephone conversations. Digital avatars offer a promising alternative, but how the brain integrates audiovisual information from such artificial sources remains unclear. Using magnetoencephalography (MEG), we investigated how the brain processes and integrates speech when visual information is provided by either natural or artificial (avatar-based) signals. Our findings reveal both shared and distinct neural mechanisms of audiovisual integration, providing critical insight into how visual input can support speech understanding in challenging listening conditions.

## Introduction

Understanding a conversational partner in acoustically challenging environments is difficult, particularly for individuals with hearing impairments [Miles et al., 2022]. Speech comprehension can be substantially enhanced when listeners can see the speaker’s face, drawing on visual information such as lip movements, facial expressions, head movements, and other subtle cues [Peelle and Sommers, 2015, Haider et al., 2022, Nirme et al., 2020, Kothe et al., 2025].

Recent advances in deep neural networks have enabled the generation of highly realistic artificial talking avatars that avoid the uncanny valley [D-ID, 2025, Song et al., 2025, Karras et al., 2017]. These avatars are increasingly used in social media and professional contexts, for example in corporate training videos, and hold considerable potential to support people with hearing impairment in everyday communication scenarios, such as public announcements in train stations [Li et al., 2024, Etibar Aliyev et al., 2025].

Several studies have demonstrated that avatars can improve speech comprehension in noisy environments [Nirme et al., 2020, Varano et al., 2022, Shan et al., 2022, Thézé et al., 2020, Cooper et al., 2025]. However, the benefit provided by avatars is consistently smaller than that achieved with natural visual stimuli. Notably, this discrepancy persists even when participants are unable to consciously distinguish avatars from natural videos [Vougioukas et al., 2020, Varano et al., 2022]. The neural mechanisms underlying the processing of talking faces generated by deep neural networks remain largely unexplored. Elucidating these mechanisms may advance our understanding of audiovisual integration and reveal neurobiological factors that currently limit the effectiveness of avatar technologies in naturalistic communication.

A key neural mechanism underlying speech perception is neural speech tracking. During speech listening, neural activity in the auditory cortex entrains to amplitude fluctuations of the speech signal [Giraud and Poeppel, 2012, Ding and Simon, 2014]. This entrainment occurs primarily in the high delta (1–4 Hz) and theta (4–8 Hz) frequency bands, which appear to serve distinct functions. Theta-band tracking is related to the syllabic rhythm of speech and reflects lower-level acoustic processing [Molinaro and Lizarazu, 2018, Etard and Reichenbach, 2019], whereas delta-band tracking corresponds to the word rate and reflects higher-level linguistic processing [Broderick et al., 2018, Weissbart et al., 2020, Gillis et al., 2021].

Neural speech tracking is also critical for audiovisual speech perception. Audiovisual speech elicits enhanced neural tracking that exceeds the superposition of auditory-only and visual-only responses [Crosse et al., 2016, Luo and Poeppel, 2007]. This enhancement is stronger in listeners with better speech-in-noise comprehension and emerges particularly under challenging acoustic conditions, an effect known as inverse effectiveness [Ross et al., 2006, Peelle and Sommers, 2015, Sumby and Pollack, 1954, Crosse et al., 2016]. Audiovisual enhancement has been observed mainly in the delta band and has not been detected above 6 Hz [Crosse et al., 2016].

Remarkably, neural tracking of speech amplitude fluctuations can occur in the auditory cortex even in the absence of sound, when only visual speech information is available [Bourguignon et al., 2020, Aller et al., 2022, Bröhl et al., 2022]. This visually-driven tracking has primarily been reported in the low delta band (0.5–1 Hz) during silent lip reading and is stronger in individuals with superior lip-reading abilities [Bröhl et al., 2022].

In the present study, we recorded magnetoencephalography (MEG) data from 32 participants who listened to continuous speech accompanied by different visual stimuli, including still images, natural videos, and avatars generated by deep neural networks (DiD AI, Israel) [D-ID, 2025]. We behaviorally assessed the participants’ speech-in-noise comprehension. We further analyzed their neural speech tracking in the low delta, high delta, and theta frequency bands to assess how different visual signals, and in particular artificial avatars, modulate audiovisual speech processing.

## Results

### Experimental Design

We recorded MEG-data while participants listened to and watched short German sentences (Fig. 1). Each sentence consisted of five words from the Oldenburger Satztest (OLSA) [Wagener et al., 1999], which were masked with four-talker babble noise. After each sentence, participants repeated every word that they understood. Some sentences were presented without the audio stimuli, and subjects were then asked to repeat the first word that they lip-read, which was always a name. We thus obtained precise behavioral data on the speech comprehension for each participant and each sentence.

**Figure 1:**
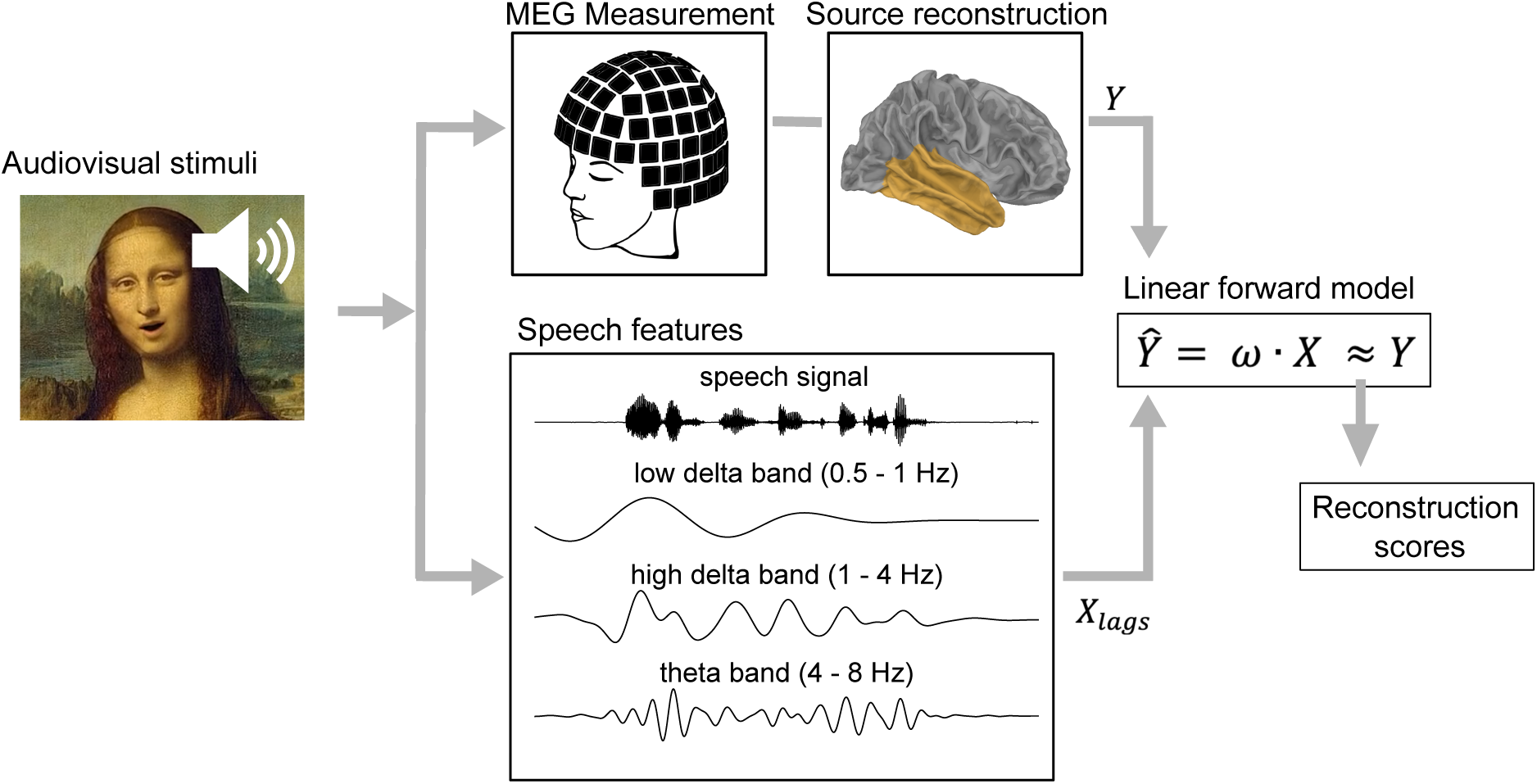
The images of the real speaker and his avatar are replaced by a speaking avatar of Mona Lisa in this Preprint. Overview of the MEG data acquisition and analysis. We created different types of audiovisual speech stimuli and presented them to 32 participants while their neural activity was recorded using MEG. The recorded MEG data were subsequently preprocessed and source-reconstructed in the auditory cortex, resulting in the neural data *Y*. The temporal envelopes of the speech stimuli were decomposed into different frequency bands, yielding the predictor variables *X*. Using a linear forward model with ridge regression, the Temporal Response Function (TRF) *w* was estimated. The TRF describes the time-lagged transformation from the stimulus features *X* to the measured neural response *Y*, yielding a predicted neural signal 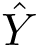. The agreement between the actual MEG response *Y* and the estimate 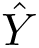 was quantified by a reconstruction score.

We employed four different types of visual stimuli (Fig. 2). These were (1) the natural video of the talker, (2) an avatar generated by a deep neural network on the basis of a still image of the talker, (3) a degraded video in which the natural video was reduced to edges of and in the talker’s face, and (4) a still image of the talker without temporal change. The degraded video and the still image thereby acted as control conditions.

**Figure 2:**
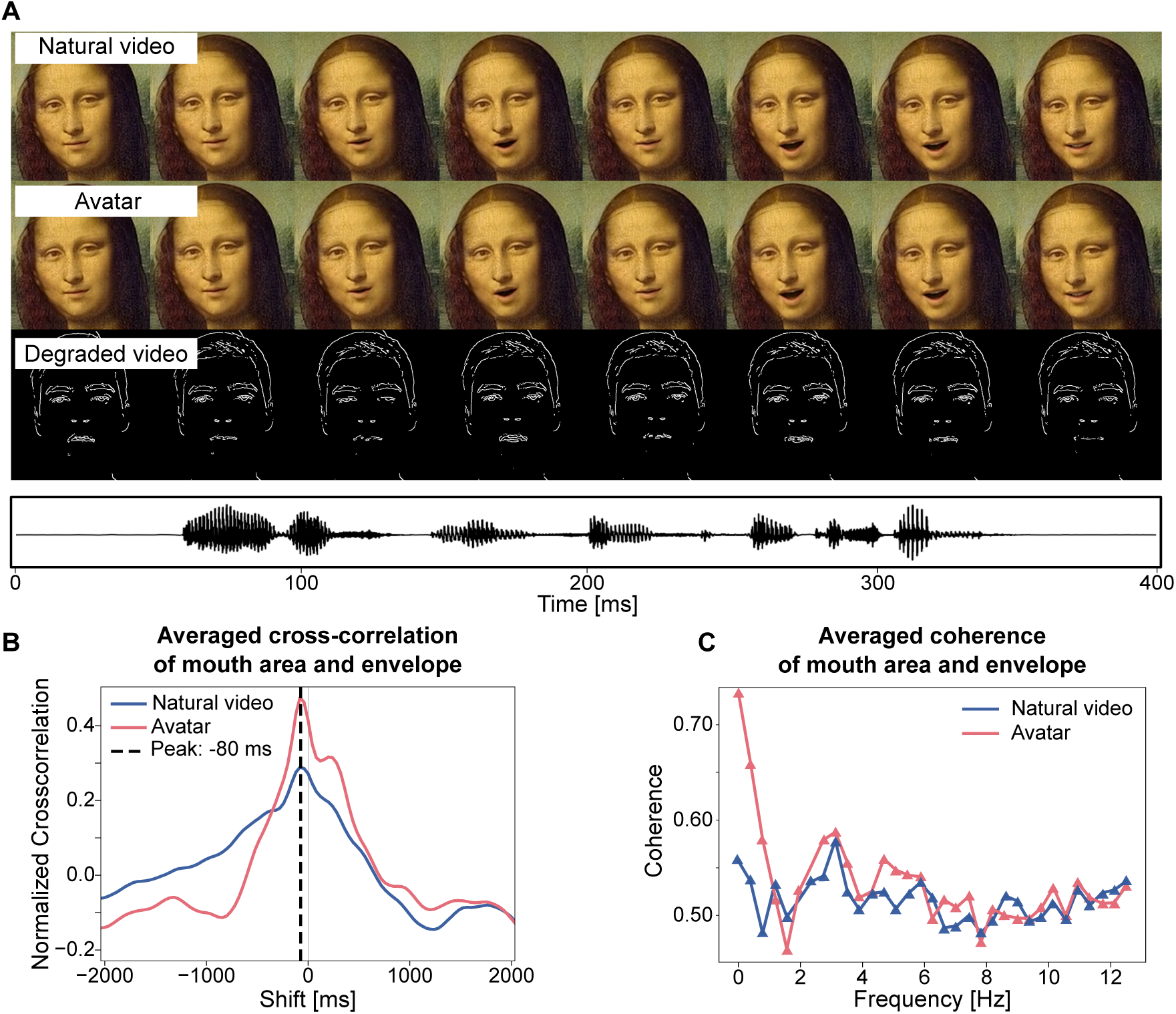
(*A*) The images of the real speaker and his avatar are replaced by a speaking avatar of Mona Lisa in this Preprint. The three animated visual stimuli are the natural video of the talker, the avatar generated from a still image of the talker through DNNs, and the degraded video, derived from the natural video but with only edges in and around the face retained. The waveform of the corresponding audio signal, a single sentence, is depicted below. (*B*) The cross-correlation between the mouth opening and the envelope of the speech signal, averaged over all employed sentences, shows a peak at-80 ms both for the natural video and the avatar. The mouth motion accordingly precedes the sound by about 80 ms. (*C*) The averaged coherence between the mouth opening area and the audio envelope, averaged across all sentences, shows comparable coherence above 1 Hz, but larger values for the avatar below 1 Hz.

The stimuli of types (1) to (3) were presented both with and without the audio stimulus. The still image in condition (4) was presented only with the audio. In the following, we refer to stimuli without the audio signal as *visual-only stimuli*. In contrast, stimuli that include the audio will be referred to as *audiovisual stimuli*. These include the speech presentation with a still image, which, due to its lack of a temporally changing visual signal, may also be viewed as an audio-only stimulus.

The MEG data were then source-reconstructed. The responses in the left and right temporal lobes were related to the speech rhythms in the low delta, the high delta, and the theta frequency bands to determine the neural speech tracking (Figure 1). To this end, we related neural activity to the speech envelope using linear forward models, yielding Temporal Response Functions (TRFs) for the three frequency ranges.

### Relation between speech envelope and mouth opening

The mouth opening of a talker is related to the envelope of the speech signal, which can play a role in audiovisual speech processing [Chandrasekaran et al., 2009]. We therefore investigated whether this relation differed between the avatars and the natural videos. To this end, we first computed the cross-correlation between the area of mouth opening of the talker and the speech envelope (Figure 2*B*). We found comparable cross-correlation functions for the two stimulus types. In particular, both peaked at the same latency of-80 ms. This showed that for both types of videos the mouth opening preceded the audio signal by 80 ms, in agreement with previous findings [Chandrasekaran et al., 2009].

We also computed the spectral coherence between the area of mouth opening and the speech envelope. Above 1 Hz, including the high delta and theta bands, coherence was similar between the avatars and the natural videos. However, below 1 Hz, in the low delta band, the avatars showed markedly higher coherence than the natural videos. The coherence at these low frequencies exceeded, for the avatars, the coherence above 1 Hz.

### Behavioral Results

We then analyzed participants’ speech comprehension across the different audiovisual stimuli. The audiovisual conditions, which included the speech signal, all had average comprehension scores above 60% (Figure 3*A*). A repeated-measures ANOVA across all audiovisual conditions revealed a significant main effect of condition on speech comprehension (*p <* 0.001). When presented with only the still image as the visual signal, participants correctly understood, on average, 66% of the words. Performance significantly increased to 78% for the avatar (*p <* 0.001). Speech comprehension of the degraded video was, on average, 76%, comparable to that for the avatar. This value was also significantly higher than that for the still image (*p <* 0.001). Given comparable levels of speech comprehension between the avatar and the degraded video, the latter could serve as a suitable control for the neural speech-tracking evoked by the avatar.

**Figure 3:**
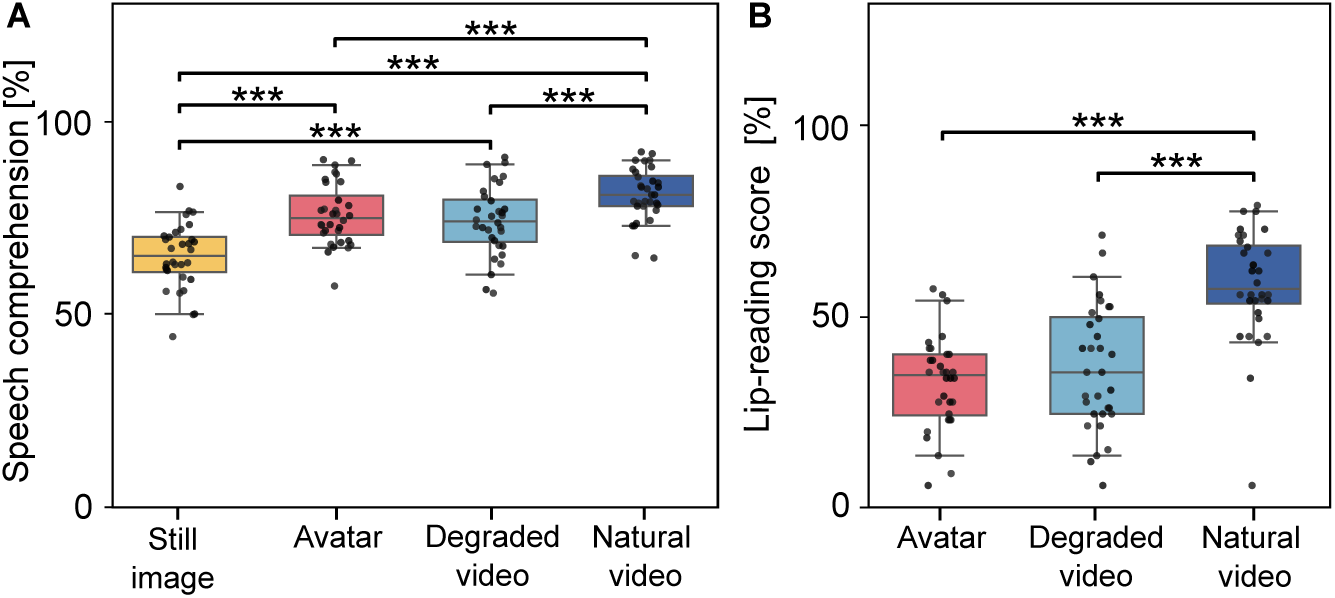
Speech comprehension scores averaged across all sentences and shown across participants for each condition. (*A*) For the audiovisual stimuli, speech comprehension was lowest in the audio-only condition (still image), increased for the avatar and the degraded video, and was highest for the natural video of the talker. (*B*), In the visual-only condition, participants could lip-read the first word of the sentence with only a low accuracy of about 35% for the avatar and the degraded video. The accuracy increased significantly to 59% for the natural video (*: *p <* 0.05; **: *p <* 0.01; ***: *p <* 0.001).

Speech comprehension was, at 84%, highest for the natural video. This value was significantly higher than the speech comprehension in the other audiovisual conditions (*p <* 0.001 for all comparisons). The *p*-values were obtained from paired t-tests with Bonferroni correction for multiple comparisons.

For the visual-only stimuli, ANOVA again showed a significant main effect across the conditions (*p <* 0.001). Participants correctly lip-read an average of 32% of the names for the avatar and 36% for the degraded video, with no significant difference between the two conditions (Figure 3*B*). These scores were above the chance level of 10% (*p <* 0.001 for the avatar and *p <* 0.001 for the degraded video). When viewing natural videos, participants performed significantly better than when stimulated with the avatar or degraded videos, correctly lip-reading 59% of the presented names (both comparisons: *p <* 0.001).

### Reconstruction scores of the linear models and relation to behavior

We constructed linear forward models to predict the MEG activity in the auditory cortex from the presented speech signal. The models yielded reconstruction scores between *−*1 and 1, which quantified the model’s performance. Here, *−*1 indicates perfect anticorrelation between the predicted neural activity and the actual neural activity, 0 indicates no agreement between the model estimate and the actual neural activity, and 1 indicates perfect reconstruction.

Linear forward models were computed for three frequency bands: low delta (0.5–1 Hz), high delta (1–4 Hz), and theta (4–8 Hz). The resulting reconstruction scores were first analysed with ANOVA. If significant differences emerged, post-hoc pairwise Student’s t-tests were conducted between the different audiovisual conditions (Figure 4*A*). To investigate potential relations between the reconstruction scores and behavior, we computed repeated-measures correlations between the scores and each participant’s average speech compre-hension across the different audiovisual conditions (Figure 4*B*).

**Figure 4:**
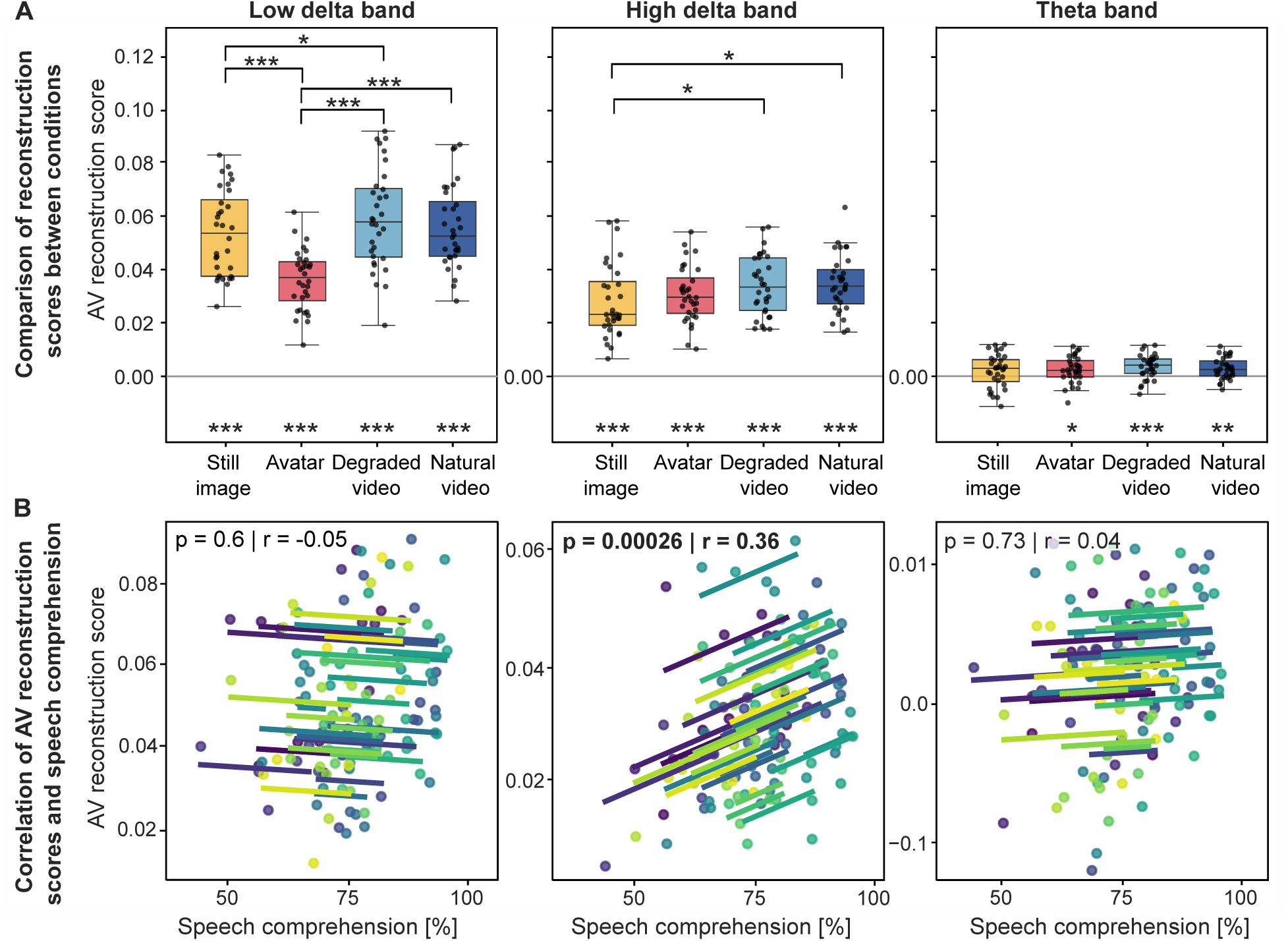
Reconstructions scores for the different audiovisual conditions and relation to speech comprehen-sion. (*A*) Reconstruction scores for the low delta band (left), the high delta band (middle), and the theta band (right). Individual points represent data from individual subjects. All scores except for the still image in the theta band were significantly larger than 0. In the low delta band, the reconstruction scores for the avatar were significantly smaller than those for the other stimuli. Delta-band reconstruction scores increased from the still image to the natural video. The scores in the theta band were much lower and did not differ significantly between the conditions. (*B*) Repeated-measures correlations between the reconstruction scores and the speech comprehension across participants. Each line corresponds to a single subject, with the corresponding data for the four audiovisual stimuli shown as data points in the same color. A highly significant positive correlation emerged for the high delta band (middle; *: *p <* 0.05; **: *p <* 0.01; ***: *p <* 0.001)

The scores in the low delta band showed clear differences between the avatar and the other audiovisual stimuli (Figure 4*A* left). In particular, ANOVA revealed significant differences between the conditions (*p <* 0.001). Subsequent pairwise comparisons showed that the scores for the still image and natural video were comparable, whereas those for the degraded video were significantly higher than those for the still image (*p* = 0.02). The highest statistical significance, however, emerged for the comparison of the avatar scores with those of the other conditions: neural tracking for the avatar was significantly weaker than that evoked by the other stimuli (*p <* 0.001 for all comparisons). No significant correlation emerged between behavioral performance and the reconstruction scores across the different audiovisual stimuli (Figure 4*B* left).

In the high delta band, the reconstruction scores also differed between the audiovisual conditions (*p* = 0.002, ANOVA). They closely followed the pattern seen in the speech comprehension scores, increasing from the still image condition to the avatar, the degraded video, and the natural video condition(Figure 4*A* middle). The differences between the scores for the still image and those of the degraded video, as well as those of the natural video, were statistically significant (*p* = 0.04 for both comparisons). The repeated-measures correlation revealed a statistically highly significant increase in the reconstruction scores, and thus in the neural speech tracking, with rising speech comprehension (*r* = 0.36, *p <* 0.001).

In the theta band, reconstruction scores were much smaller than in the low and high delta bands, and even negative for some subjects (Figure 4*A* right). Statistical testing revealed that, on the population level, the scores for the avatar, the degraded video, and the natural video still significantly exceeded the baseline value of 0. For the still image, i.e., in the audio-only condition, however, the scores were not statistically different from 0. ANOVA did not show statistically significant differences between the conditions (*p* = 0.06). No statistically significant correlation was found between the reconstruction scores and speech comprehension (Figure 4*B* right).

The repeated measures correlation between neural tracking and speech comprehension ensured that within-condition variability was not driving between-condition effects. In a second analysis, we also tested whether, within each condition and each frequency band, the reconstruction of the participants’ MEG data correlated with their word comprehension (Figure 5*A*). We found statistically significant positive relations in the low delta band for the avatar (Figure 5*A* left) and in the delta band for the natural video (*r* = 0.5, *p* = 0.01 resp. *r* = 0.5, *p* = 0.01, Figure 5*A* left and middle). The theta band did not yield a significant correlation (Figure 5*A* right).

**Figure 5:**
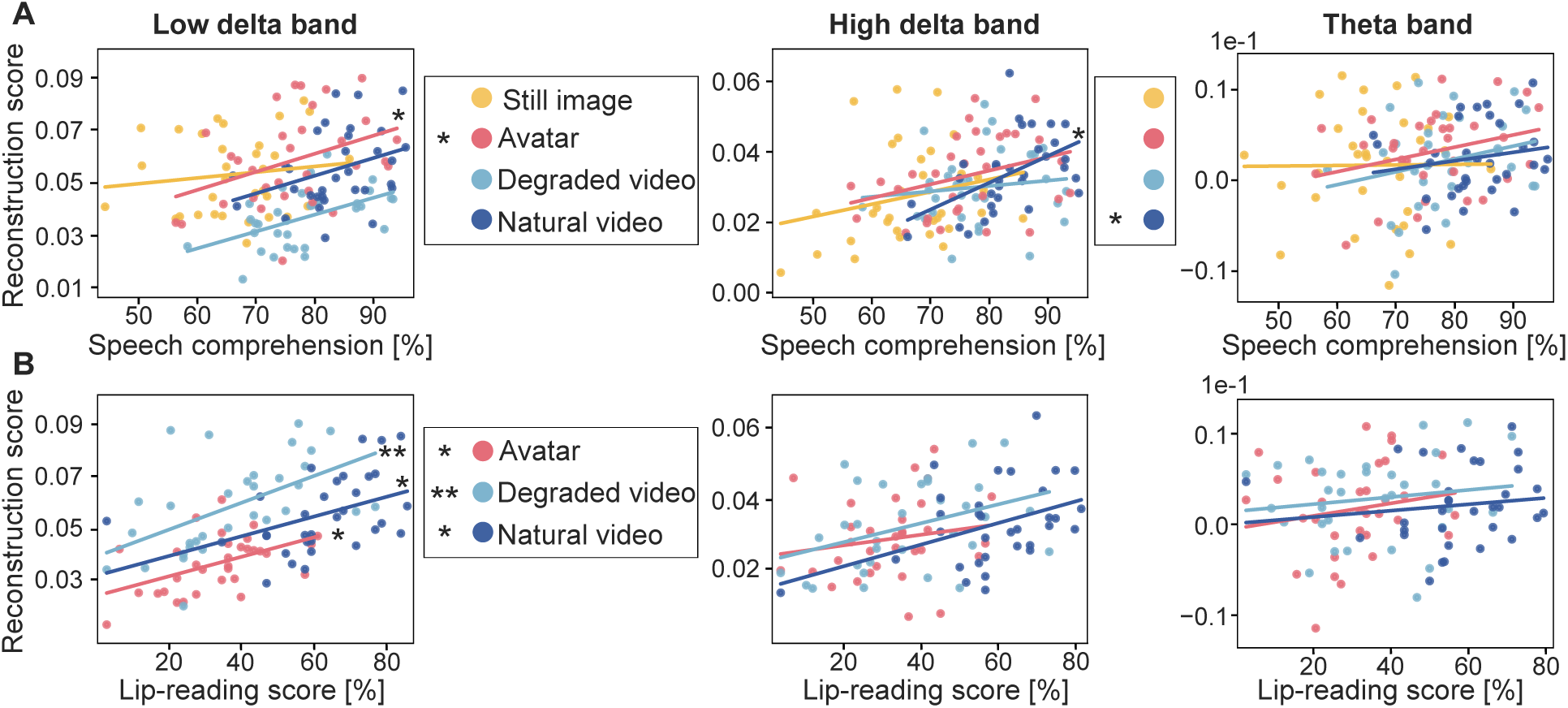
Within-condition correlations between reconstruction scores and behavioral scores. (*A*) Within-condition correlations between the reconstruction scores and speech comprehension for the three frequency bands. Significant correlations emerged for the avatar in the low delta band and for the natural video in the high delta band. (*B*) Within-condition correlations between the reconstruction scores and the lip-reading performance for the visual-only stimuli. All audiovisual conditions in the low delta band, but none in the high delta and theta bands, were significant (*: *p <* 0.05; **: *p <* 0.01; ***: *p <* 0.001).

Because the neural response to the audiovisual speech stimuli could be (partly) driven by the lip-reading skills of the participants, we additionally correlated the reconstruction scores of the MEG responses to the audiovisual stimuli with the lip-reading scores obtained in the visual-only conditions (Figure 5*B*). These correlations were insignificant in the high delta and in the theta band (Figure 5*B*). However, in the low delta band, we found statistically significant positive correlations for the avatar (*r* = 0.5, *p* = 0.01), for the degraded video (*r* = 0.5, *p* = 0.005), and for the natural video (*r* = 0.4, *p* = 0.04). All within-condition correlations were computed using Pearson’s correlation coefficient, and the *p*-values were corrected for multiple comparisons with the Bonferroni method.

### Temporal Response Functions (TRFs)

The temporal response functions (TRFs) provide further information on the neural speech tracking. In particular, they exhibit peaks at particular temporal delays, highlighting the latencies at which the neural tracking arises (Figure 6). As an example, in the low delta band, a prominent peak occurs at a delay of about 240 ms (Figure 6*A* left). Due to the similarity to the event-related potential measured with MEG at a delay of 200 ms, we refer to this peak as the *M* 200_TRF_.

**Figure 6:**
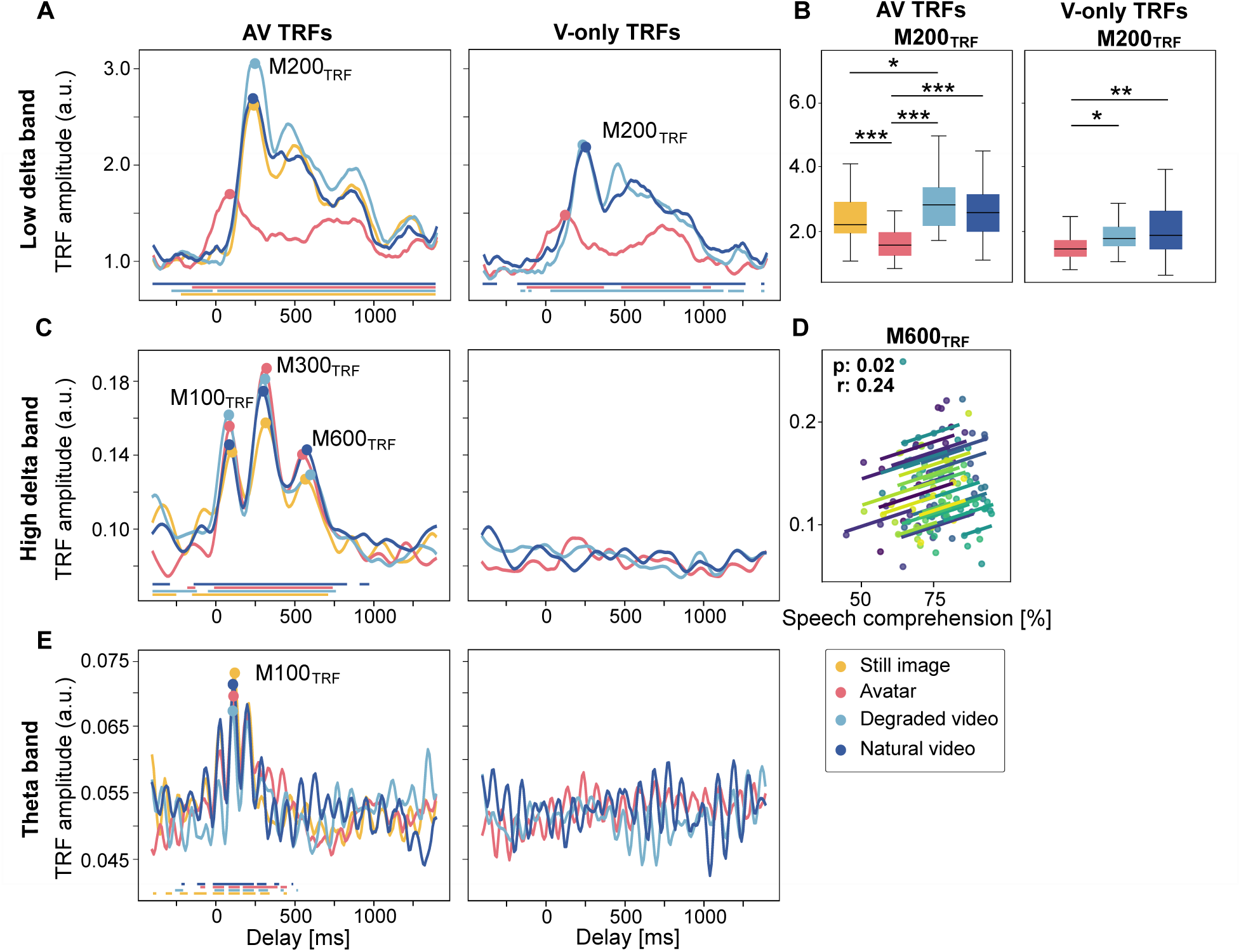
Temporal Response Functions (TRFs). (*A*) The TRFs in the low delta band exhibited a significant peak at a latency of about 240 ms (*M* 200_TRF_). The delay was earlier for the avatar compared to the other stimuli. The peaks appeared both for the audiovisual (right) and for the visual-only stimuli (left). (*B*) The amplitude of the peak *M* 200_TRF_ was significantly smaller for the avatar than for the other stimuli, both for audiovisual (left) and for visual-only (right) stimulation. (*C*) In the audiovisual conditions (left), the high-delta band TRFs displayed the pronounced peaks *M* 100_TRF_, *M* 300_TRF_, and *M* 600_TRF_. No such peaks emerged for visual-only stimulation (right). (*D*) Repeated measures correlation revealed a significant relation between the amplitude of the peak *M* 600_TRF_ and speech comprehension. Each color denotes an individual participant. (*E*) The theta-band TRFs displayed peaks for the audiovisual stimuli (left) but not for the visual-only conditions (right). Comparisons between stimulus conditions were assessed using paired t-tests, and the Bonferroni-corrected significance levels are indicated by asterisks (*: *p <* 0.05; **: *p <* 0.01; ***: *p <* 0.001).

We computed TRFs in the low delta, high delta, and theta bands. Because silent lip reading of videos without audio has been shown to induce neural tracking of speech rhythms in the low delta band, we also computed TRFs for the visual-only stimuli [Bourguignon et al., 2020, Aller et al., 2022, Bröhl et al., 2022].

To examine if the magnitude of a peak in a TRF differed between the audiovisual stimuli, we employed ANOVA. If this test yielded a significant overall effect in a certain frequency band, we compared the magnitudes of the peak pairwise across the different audiovisual conditions in this frequency band. Only the low delta band yielded significant differences in ANOVA, so we only show the pairwise comparisons for this frequency band (Figure 6 in part *B*).

Additionally, for each identified TRF-peak, an across-condition analysis of the relationship between peak amplitude and speech comprehension was performed. To this end, we computed a repeated measures correlation between the amplitude of the TRF peak and each participant’s comprehension scores across all four audiovisual conditions. Again, only the data for the frequency bands with significant results are presented in Figure 6, resulting in a single plot for the high delta band (Figure 6*D*, each color denotes an individual subject).

Similar to the reconstruction scores, in the low delta band, the TRFs for the avatar differed significantly from all other stimuli. In particular, the TRFs for the avatar showed a much earlier neural entrainment with a peak at 90 ms compared to the other conditions (still image: 230 ms; degraded and natural video: 250 ms, (Figure 6*A* left). Moreover, ANOVA showed significant differences in peak amplitudes across conditions (*p <* 0.001). Post-hoc testing revealed that the peak amplitude for the avatar was significantly smaller than those in the other conditions (*p <* 0.001 compared to the still image, the degraded video, and the natural video; Figure 6*B* left). As seen in the reconstruction scores, we also observed a small but significant difference in the amplitude of the TRF peak between the degraded video and the still image (*p* = 0.01)Figure 6*B* left. A repeated measures correlation between the peak amplitude and the comprehension scores did not yield a statistically significant result, and the corresponding plot is therefore not presented.

The TRFs obtained for the visual-only stimuli in the low delta band also showed a neural response in the auditory cortex (Figure 6*A* right). The TRFs were similar to the ones obtained for the corresponding audiovisual condition but had a lower amplitude. The peak amplitude varied significantly between the different conditions (*p <* 0.001, ANOVA). The amplitude of the peak for the avatar was again smaller than that for the degraded video (*p* = 0.02) and that for the natural video (*p* = 0.004, Figure 6*B* right). There was no significant correlation between the peak amplitude and the lip-reading scores.

The TRFs in the high delta band revealed three peaks at delays of 90 ms, 300 ms, and 570 ms that we refer to as *M* 100_TRF_, *M* 300_TRF_, and *M* 600_TRF_, respectively (Figure 6*C* left). The variations of these peak amplitudes across the different stimuli were, however, not statistically significant (*p* = 0.23, *p* = 0.19, and *p* = 0.40, ANOVA). However, the amplitude of the peak *M* 600_TRF_ increased in line with increasing speech comprehension. For this peak, we obtained a significant correlation between the peak amplitude and speech comprehension across the audiovisual conditions (*r* = 0.24, *p* = 0.02, Figure 6*D*).

The TRFs for the visual-only conditions in the high delta band showed no neural entrainment in the auditory cortex (Figure 6*C* right).

The theta-band TRFs obtained from the audiovisual stimuli exhibited multiple peaks (Figure 6*E* left). The largest peak occurred at a latency of 110 ms and is referred to as *M* 100_TRF_. Due to the narrow-band nature of the theta frequency band, the neighboring peaks are likely sidelobes and originate from the same neural source. We therefore focused our analysis on the peak *M* 100_TRF_. Its amplitude did, however, not vary significantly between the audiovisual conditions (*p* = 0.66, ANOVA). The repeated-measures correlation between peak amplitude and speech comprehension was not statistically significant either.

As in the high delta band, the theta-band TRFs did not show significant responses in the visual-only conditions (Figure 6*E* right).

## Discussion

We investigated how artificially created talking avatars influence the neural tracking of speech rhythms in the auditory cortex. Participants were presented with five-word matrix sentences accompanied by a still image of the talker, an avatar, a degraded version of the natural video, or the natural video itself. Visual-only conditions without accompanying audio were also included. Behaviorally, speech comprehension improved systematically with increasing visual information, from the audio-only condition to the avatar and degraded video, and further to the natural video.

This behavioral pattern was reflected in neural speech tracking in the high delta frequency band. In contrast, theta-band responses did not differ across audiovisual stimuli. Strikingly, neural speech tracking in the low delta band was strongly altered for the avatar condition, showing reduced magnitude and markedly earlier latency compared to all other stimuli.

### Speech comprehension

The behavioral data followed a plausible pattern: participants’ performance improved with increasing availability of visual information. The reduced speech comprehension for the degraded video, as compared to the natural one, probably resulted from the reduced visual information that nonetheless retained accurate lip movements.

Behavioral performance increased with the availability and quality of visual information. Reduced speech comprehension for the degraded video compared to the natural video likely resulted from diminished visual detail, despite preserved lip movements. The audiovisual benefit of the avatar fell between that of the audio-only condition and the natural video, consistent with previous studies showing that avatars can support speech comprehension, albeit less effectively than natural facial videos and depending on the naturalness of facial dynamics, lip synchrony, and head movements [Munhall et al., 2004, Varano et al., 2022, Shan et al., 2022].

In visual-only conditions, lip-reading performance was highest for the natural video and nearly matched performance in the audio-only condition. This likely reflects the task design: for the visual-only stimuli, participants repeated only the first word (a name) of each sentence. Given the limited set of possible names in the OLSA matrix test, participants could learn to identify them from minimal visual cues. This design was chosen to avoid participant frustration that could arise from full-sentence lip reading.

Importantly, visual-only avatar and degraded video conditions yielded comparable performance above chance level. Together with the audiovisual results, this suggests that the degraded video provided a behavioral benefit similar to the avatar for both speech comprehension and lip reading, justifying its use as a control stimulus in the neural analyses.

### Neural speech tracking in the high delta band predicts audiovisual speech comprehension

Neural tracking in the high delta band (1 *−* 4 Hz) increased from the audio-only condition to the animated stimuli (avatar, degraded video, and natural video), paralleling the increase in speech comprehension scores. Across audiovisual conditions, the strength of neural tracking correlated strongly with behavioral comprehension. For natural video stimuli, participants with higher comprehension scores exhibited stronger high-delta tracking, consistent with previous findings [Aller et al., 2022, Luo and Poeppel, 2007]. In contrast, visual-only stimuli did not elicit high-delta tracking, despite successful lip reading, in line with earlier MEG studies reporting no auditory cortical tracking above 1 Hz in the absence of sound [Bourguignon et al., 2020].

These results support the role of high-delta-band tracking as a neural marker of audiovisual integration. Artificially generated visual signals can enhance speech understanding by boosting neural tracking in this frequency band, consistent with earlier work linking low-frequency cortical phase locking and coherence to speech comprehension [Luo and Poeppel, 2007, Crosse et al., 2016, Aller et al., 2022] and with evidence that audiovisual speech improves the accuracy of cortical speech tracking [Peelle and Sommers, 2015, Crosse et al., 2015].

TRF analyses revealed that audiovisual integration primarily occurred at a late latency of approximately 600 ms. The amplitude of this *M* 600_TRF_ component correlated significantly with speech comprehension. This late response resembles the P600 component of event-related potentials, which has been associated with semantic and syntactic processing as well as lexical retrieval during spoken language comprehension [Bornkessel-Schlesewsky and Schlesewsky, 2008, Gouvea et al., 2010, Brouwer et al., 2012].

To our knowledge, this study is the first to link the *M* 600_TRF_ of neural speech tracking to audiovisual speech processing. Neural tracking in the delta band aligns with the rhythm of words in speech and has previously been associated with linguistic processing [Broderick et al., 2018, Weissbart et al., 2020, Etard and Reichenbach, 2019].

In contrast, earlier speech tracking components at latencies around 100 ms and 300 ms were not related to comprehension, suggesting that they reflect lower-level acoustic processing largely unaffected by higher-level linguistic factors.

### Neural tracking in the low delta band is impaired when viewing the avatar

All audiovisual stimuli elicited robust speech tracking in the auditory cortex in the low delta band (0.5 *−* 1 Hz). Notably, this was the only frequency band that showed neural speech tracking in the absence of sound, that is, during visual-only conditions. This finding is consistent with previous reports of auditory cortical tracking during silent lip reading restricted to frequencies below 1 Hz [Bourguignon et al., 2020, Bröhl et al., 2022].

Crucially, low-delta tracking differed substantially for the avatar compared to all other stimuli. For the still image, degraded video, and natural video, responses peaked at a latency of approximately 240 ms. In contrast, the avatar elicited markedly earlier responses (around 90 ms) with roughly half the magnitude. These differences were present for both audiovisual and visual-only conditions.

This dissociation suggests that very slow cortical dynamics below 1 Hz are particularly sensitive to incon-sistencies between visual and auditory cues. Whereas the natural and degraded videos provided temporally congruent audiovisual information, and the still image provided no dynamic visual input, the avatar likely introduced subtle mismatches in visual–acoustic timing that disrupted neural tracking at these low frequencies.

Supporting this interpretation, coherence analysis between mouth movements and the speech envelope re-vealed that the avatar’s lip movements were more tightly synchronized to speech amplitude below 1 Hz than those of the natural video. For real speakers, coherence between mouth opening and the speech envelope typically peaks at higher frequencies (3 *−* 4 Hz) [Pedersen et al., 2022]. Although stronger low-frequency synchrony might appear beneficial, it likely reflects deviations from natural anticipatory mouth movements, thereby altering visual dynamics and impairing the neural readout of low-frequency speech information.

Low-delta speech tracking was present even without sound, indicating that it partly reflects lip reading. This interpretation was supported by the finding that the tracking strength in this band correlated with individual lip-reading abilities, an effect not observed in higher frequency bands. In contrast, when relating the tracking strength to speech comprehension in the presence of audio, a significant correlation emerged only for the avatar condition. Together, these results underscore the specific role of low-delta speech tracking in lip reading [Bourguignon et al., 2020, Bröhl et al., 2022] and further highlight its dissociation for avatar-based stimuli.

Because low-delta frequencies fall below the typical word rate of speech, they may be associated with pro-cessing phrasal units consisting of multiple words. The mechanisms by which speech tracking at this phrasal rate supports lip reading, as well as the specific causes of avatar-related deviations at these frequencies, remain important topics for future research.

### Neural tracking in the theta band is not informative on audiovisual speech comprehension

Speech tracking in the theta band (4 *−* 8 Hz) did not show audiovisual integration for any of the audiovisual conditions. This finding aligns with prior studies reporting audiovisual integration primarily in lower frequency bands, such as 2 *−* 6 Hz [Crosse et al., 2015]. No differences were observed between the avatar and other visual stimuli, and no theta-band tracking was detected in the auditory cortex during visual-only conditions.

The theta band corresponds to the syllabic rhythm of speech and has been proposed to support segmentation of the speech stream into syllabic units. Our results suggest that audiovisual integration does not substantially modulate this lower-level acoustic processing stage in the auditory cortex.

## Conclusions

Our study shows that artificially generated talking faces can enhance speech-in-noise comprehension, even though they elicit impaired neural tracking in the low delta band. At the same time, our results corroborate the crucial role of the low delta band in silent lip reading: neural entrainment to the speech envelope emerged in the auditory cortex, specifically in the low delta band, but not in other frequency bands. Moreover, low-delta cortical tracking was significantly correlated with participants’ lip-reading abilities.

These findings allow for two possible interpretations. First, low-delta cortical tracking may play no role in audiovisual speech processing and may instead be specific to silent lip reading. On this account, audiovisual speech processing and lip reading would rely on entirely distinct neural mechanisms.

Second, it is possible that low-delta tracking does contribute to audiovisual speech processing and that its impairment when viewing the avatar limited the avatar’s effectiveness in improving speech-in-noise comprehension. If so, improving the avatar so that it more faithfully represents mouth movements at frequencies below 1 Hz should restore both low-delta speech tracking and the speech-in-noise benefit. Indeed, although the avatar used in our study did enhance speech comprehension, it did not match the benefit obtained with natural stimuli. Our findings may therefore point toward a path for further increasing the speech-in-noise benefits of artificially generated talking faces.

## Materials and methods

### Audiovisual stimuli

We employed four different audiovisual stimuli: a still image of the talker (audio-only condition), an artificially generated avatar, a degraded video of the talker, and a natural video of the talker.

The natural videos were obtained from a native German speaker uttering short five-word matrix sentences, randomly selected from the Oldenburger Satztest (OLSA) [Wagener et al., 1999]. These syntactically correct but semantically meaningless sentences consist of a name followed by a verb, a number, an adjective, and an object. There are ten possibilities for the words at each of the five positions in the sentence.

Videos were recorded at a sampling rate of *29.97* fps and an audio sampling rate of *44.1* kHz. After manually checking all 600 recorded sentences for errors such as mispronunciation or background noise, we obtained recordings of 571 sentences. These were then processed into the different audiovisual and visual-only conditions. Example frames are shown in Figure 2.

The degraded videos were obtained from the natural videos using FFmpeg’s *edgedetect* filter, which extracts and renders prominent contours while suppressing non-edge information.

To generate the avatar, we employed software from the company D-iD, which utilized a CNN-based image encoder to process a still image of the talker and a GAN image-to-video model to animate lip movements in synchrony with the input audio [D-ID, 2025].

The sentences were mixed with four-talker babble noise [Ma et al., 2009] starting at varying timings before the sentence onset. The signal-to-noise ratio (SNR) was determined in a pre-study aimed at achieving 50% speech comprehension in the audio-only condition. This resulted in an average SNR of about *−*4 dB, which was then applied for all participants in the main study.

Three types of visual-only stimuli were obtained from the avatar, the degraded video, and the natural video by muting the audio channels.

Example frames occurring every 50 ms of the natural video, the avatar, and the degraded video are shown in Figure 2.

### Area of Mouth Opening

The area of mouth opening was extracted from each video frame using the OpenFace toolkit [Baltrušaitis et al., 2016], which provides frame-wise facial landmark detection. The enclosed area of the inner lip landmarks was used as a continuous measure of mouth opening over time. We computed the cross-correlations of the resulting time series with the speech envelope, as well as the coherence between the two measures using all 571 sentences, with lags between *±*2000 ms (Figure 2B).

### Experimental design

Thirty-three healthy, right-handed participants (17 female; age 19–31 years) took part in two experimental sessions, each of which lasted up to 60 minutes, and which were spaced one to three weeks apart. One participant did not finalize the study, such that data from 32 participants were analyzed.

To determine each participant’s SRT_50_, subjects completed 80 audio-only training trials at the start of the first session with adaptive noise adjustment. This also served to familiarize the subjects with the limited word pool of the matrix test and to reduce subsequent learning effects.

Participants were then presented with the four audiovisual stimuli and the three visual-only stimuli. The presentation alternated between blocks of audiovisual stimuli and blocks of visual-only stimuli. Each block contained three sentences from each condition, which were selected randomly. The order of presentation was also random, and sentences were not clustered in conditions. Sixty sentences were presented for each stimulus type, yielding 20 audiovisual and 20 visual-only blocks. Eight blocks of each type were presented in the first session and 12 in the second. Each sentence lasted about 3 s, yielding about 3 minutes of data per condition.

After each audiovisual sentence, participants were asked to repeat all words they understood while viewing the OLSA word matrix. After a visual-only sentence, they were asked to articulate the name, i.e., to lip-read the first word in the sentence.

### Data acquisition and preprocessing

MEG data were recorded with a 248-channel whole-head system (4D-Neuroimaging) at 1, 017.25 Hz. Par-ticipants lay in a supine position with eyes open in a magnetically shielded chamber. Online data preprocessing included analog band-pass filtering between 1–200 Hz. In addition, the system used a calibrated linear weighting of 23 reference sensors developed by 4D Neuroimaging (San Diego, CA, USA). These sensors enabled effective correction of environmental noise, thereby improving the accuracy and reliability of the recorded data. The filter boundaries were not sharp, such that activity in the low delta band (0.5–1 Hz) remained. The head shape and five fiducial landmarks were digitized using a Polhemus system before each session.

Offline preprocessing was performed in MNE-Python [Gramfort et al., 2014]. Power-line noise was removed with a 50 Hz notch filter (FIR, transition bandwidth 0.5 Hz), and the data were resampled to 100 Hz.

Auditory stimuli were delivered via a custom-built system [Schilling et al., 2020], with precise temporal alignment ensured by recording the stimulus signal through an MEG input channel and synchronizing it via cross-correlation with the actual audio, achieving accuracy up to 1 ms. Visual stimuli were projected onto a screen above the participant’s head. A constant audio–visual delay introduced by the presentation system was measured and corrected for [Varano et al., 2023].

### Neural Source Localization

Source reconstruction was performed in MNE-Python [Gramfort et al., 2014] using the FreeSurfer “fsaverage” template [Fischl, 2012]. Individual head positions and digitized head shapes were used to co-register the template via rotation, translation, and scaling.

A 5 mm volumetric source grid was created, and cortical ROIs were defined based on aparc parcellation, including the bilateral auditory cortical regions (inferior temporal gyri, middle temporal gyri, superior temporal gyri, transverse temporal gyri, banks of the superior temporal sulci, entorhinal cortices, parahippocampal gyri, fusiform gyri, and temporal poles) as these have been shown to be sensitive to audiovisual integration [Van Atteveldt et al., 2010, Macaluso et al., 2004]. This yielded 862 voxels. A boundary element model (BEM) was used for forward computation. Source activity was estimated with a linearly constrained minimum variance (LCMV) beamformer [Bourgeois and Minker, 2009], using empirical and noise covari-ance matrices from MEG and empty-room data, respectively. Filters were normalized for unit noise gain and their orientation constrained to cortical surface normals.

### Behavioral Analysis

Participants repeated every word they understood after each presented AV sentence and every name they lip-read after each V-only sentence. From this data, average speech comprehension and lip-reading scores were calculated for each participant and stimulus.

### Linear forward models to determine neural speech tracking

Linear forward models were computed to predict source-localized neural activity from fluctuations in speech amplitude and to assess neural speech tracking. The speech envelope was obtained from the audio signal through a Hilbert transform. The envelope was then band-pass filtered in three frequency ranges, the low delta band (0.5–1 Hz), the high delta band (1–4 Hz), and the theta band (4–8 Hz). These three band-pass filtered signals were used as predictors in the models.

The linear mapping between the *i*th predictor *x_i_*(*t*) at time *t* and the MEG response *y*^(*j*)^(*t*) at voxel *j* and time *t* was formulated using weights at different time delays [Lalor and Foxe, 2009]:

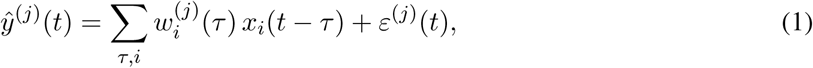

in which 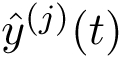 denotes the predicted neural response at voxel *j* and time *t*, *x_i_*(*t* − *τ*) is the *i*th stimulus feature at time lag *τ*, the weight *w_(i)_*^(^*^j^*^)^(*τ*) represents the TRF for the *i*th predictor and the neural activity at voxel *j* at time lag *τ*, and *ε_(i)_*^(^*^j^*^)^(*t*) is the residual error term. The model was trained using ridge regression, which minimizes the following cost function:

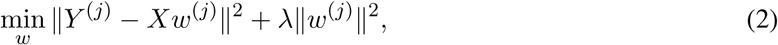

in which *X* is the matrix of time-lagged predictors, *Y* ^(*j*)^ the measured MEG signal at voxel *j*, and *λ* the regularization parameter.

Stimulus and MEG data were z-scored and concatenated per condition and participant. Ridge regression was conducted with five-fold cross-validation across the parameter range *λ* = [10*^−^*^7^,…, 10^7^].

Pilot analysis showed that responses in the low delta band were stronger than those in the higher frequency bands. These responses bled into the high delta band and required a lower value of *λ* for the optimal fit than the other frequency bands. To prevent cross-frequency bleeding, we first computed a forward model for neural tracking in the low delta band. This yielded an optimal value of *λ* = 1. We then subtracted the predicted neural responses from the actual ones. Using the residual neural data, we then computed forward models for the speech envelope filtered in the high-delta and theta bands. We obtained a higher optimal value of *λ* = 100.

The TRFs were computed at a sampling frequency of 100 Hz, yielding 200 lags between *−*500 ms and 1, 500 ms for each of the 862 voxels. Reconstruction accuracy was quantified by the Pearson correlation coefficient between the predicted and the measured signals, averaged across all voxels. The accuracy was determined from the testing fold of the five-fold cross-validation.

Population-level TRFs were obtained by averaging the absolute values across voxels and participants. The latencies of the peak responses were identified in the averaged TRF. The latencies of the response peaks for the individual participants were determined within *±*50 ms of the population-averaged latencies and were extracted for statistical analysis.

## Statistical Analysis

To test whether reconstruction scores of the linear forward models and the TRF amplitudes were significantly different from noise, the reconstruction scores were compared against zero, and the TRFs were tested against a null model generated from randomly mismatched pairings of MEG segments and audio sentences. This random mismatching was conducted five times per participant, resulting in 160 null-model TRFs across the 32 participants. These 160 null models were the basis for generating a null distribution of 500 bootstrapped null models. We then calculated empirical *p*-values for each time lag for the population-level TRF.

To compare behavioral or neural metrics across the different types of stimuli, we first conducted an ANOVA to determine whether there were significant differences. If so, we then performed pairwise comparisons using paired-sample *t*-tests with Bonferroni correction for multiple comparisons. The relationships between the reconstruction scores of the linear models and behavioral performance across the different conditions were analyzed using repeated-measures correlations and Pearson’s correlation coefficient, with Bonferroni correction for multiple comparisons.

## Data and Code Accessibility

The data and code used in this study are available from the authors upon request.

## Acknowledgments

This project was supported by the German Science Foundation (DFG, project number 523344822).

## Author contributions

JR and TR designed research; JR performed research; JR and AS collected the data; JR and TR analyzed data; JR wrote the paper; AW, SZ, DK, TR, and CJ interpreted the results and provided critical feedback.

